# Assessing Human Liver Spheroids as a Model for Antiviral Drug Evaluation Against BSL-3 Haemorrhagic Fever viruses

**DOI:** 10.1101/2025.01.29.635416

**Authors:** Sarah Chaput, Jean-Sélim Driouich, Xavier de Lamballerie, Antoine Nougairède, Franck Touret

## Abstract

Haemorrhagic fever viruses (HFVs) cause highly lethal syndromes with limited therapeutic options. Increasingly, 3D cell culture models are becoming an important tool in the field of virology. Since the liver is an important target for many HFVs, we evaluated a ready-to-use 96-well liver spheroid model composed of human primary cells for antiviral assessment. We worked with four biosafety level 3 (BSL-3) HFVs in this study: two orthoflaviviruses, Alkhumra haemorrhagic fever virus (AHFV) and yellow fever virus (YFV), and two viruses belonging to *Hareavirales* order, Pirital virus (PIRV), a surrogate for new-world BSL-4 mammarenaviruses, and Rift Valley fever virus (RVFV). We found that RVFV and PIRV were able of replicating in this model, whereas the orthoflaviviruses were not. A high viral dose was required for robust replication, and infectivity of RVFV in spheroids was low. We successfully demonstrated the antiviral activity of known broad-spectrum antiviral compounds— favipiravir, nitazoxanide, ribavirin, and galidesivir—despite some variability. However, except for ribavirin, higher doses were required in spheroids to detect antiviral effect compared to the 2D cell culture model. Overall, we conclude that while this model can provide valuable complementary information to traditional models, it should not be considered a critical selection step for antiviral compounds. More broadly, this model could be useful to study viral pathogenicity and host-pathogen interactions.

## 1. Introduction

Haemorrhagic fever is a severe complication of viral infection caused by viruses belonging to different families (*Nairo-, Phenui-, Hanta-, Arena-, Flavi-, Filoviridae*) ^1,2^. All these viruses share common features such as an envelope and a single stranded RNA genome. Most of them are zoonotic with diverse reservoirs like domestic animals, bats, arthropods and rodents ^3^. To date, haemorrhagic fever viruses (HFVs) have been reported in Africa, America, Europe and Asia ^1,4^. HFVs use a variety of transmission routes to infect humans, including the bite of an arthropod vector, direct contact with infected faeces, or inhalation of aerosols ^1^. Human-to-human transmission may also occur with certain HFVs ^1,5^.

Once infected, the risk of developing haemorrhagic fever varies according to the virus ^1,4^. Key symptoms include fever, vascular regulation disruption, various degrees of haemorrhage and organ failure ^3,4^. Dendritic cells and macrophages are usually the first target of these viruses causing an increase in cytokine release ^6^. These cells then assist in systemic dissemination to organs, including the liver ^4,7^. The degree of liver involvement varies between viruses from mild to extensive damage, the most severe resulting from an infection with Crimean Congo haemorrhagic fever virus, Rift Valley fever virus (RVFV) or yellow fever virus (YFV) ^6^. Many HFVs infect and induce hyperplasia of Kupffer cells and necrosis or apoptosis of hepatocytes ^6^. This induce cytolytic hepatitis, often accompanied by hepatocellular insufficiency, leading to reduced production of clotting factors ^4,6^.

Currently, there is a lack of therapeutic solutions for almost all HFVs ^3^ and many HFVs are listed by the World Health Organization as priority pathogens for research and development in public health emergency contexts ^8^. Antiviral drug development classically begins with the evaluation of antiviral activity *in vitro* using 2D cell culture models, followed by validation *in vivo* with animal models ^9,10^. *In vitro* 2D cell models, typically using immortalised cell lines, are simple and cost-effective models that are relatively easy to implement in high-containment laboratories. However, although they represent a useful model, especially during initial screening stages of antiviral development, these models, often genetically altered, have the inconvenience of lacking the functions and interactions needed to faithfully mimic the biological reality of the human body ^9^. This is responsible for several failures in transition from *in vitro* to *in vivo* studies ^10,11^, costing a lot of time and money, especially when animal studies have to be performed in biosafety level 3 (BSL-3) and 4 laboratories. Therefore, it is of interest to develop more complex and predictive models to fill the gap between *in vitro* 2D cell culture models and *in vivo* animal models ^9–11^. Such models would also allow to decrease the number of animals used in accordance with the 3R rule (replacement, reduction and refinement) 11,12.

In this way, 3D culture models including spheroids, organoids, organs-on-chip and explant cultures, are strong candidates that have benefited significant development in recent years ^9,10^. Furthermore, the SARS-CoV-2 pandemic and the subsequent relentless search for an antiviral treatment highlighted the value of this modelling approach. This led to the example of chloroquine, which, following the demonstration of its *in vitro* activity in 2D models, was tested in numerous clinical trials despite its total lack of activity in primary human bronchial cells or in animal models ^13,14^. However, the main limitation of these models for drug development is the ability to use them in high throughput screening. For this purpose, spheroids are the easiest to scale today ^9,11^, with cultures from 96 to 1536-well plates. Spheroids are 3D-self aggregated cells cultivated with or without scaffold like hydrogel or matrigel ^9^. They can consist of differentiated or undifferentiated cells, with multiple cell types, and are organised into compact structures that recapitulate cell-to-cell and cell-extracellular microenvironment interactions. Scaffold-free spheroids even produce their own extracellular matrices, avoiding interferences with extern scaffold ^9^.

Since the liver is an important target for many HFVs, we evaluated in this study a human liver spheroid (HLS) model. These spheroids are scaffold-free and composed of around 1,000 primary cells including multi-donor human hepatocytes and non-parenchymal cells (NPC; liver endothelial cells and Kupffer cells) which are frequently the target of HFVs ^6^ (Fig. 1A). These organoids are ready-to-use in an appropriated 96-well format suitable for high-throughput screening of antiviral molecules. We first evaluated its permissiveness to several BSL-3 HFVs: two orthoflaviviruses, Alkhumra haemorrhagic fever virus (AHFV) and YFV, and two viruses belonging to *Hareavirales* order (formerly Bunyavirales) ^2^, Pirital virus (PIRV), as a surrogate for new-world BSL-4 mammarenaviruses, and RVFV. Then, we evaluated the efficacy of well-characterized antiviral compounds, comparing the results to data from a conventional 2D *in vitro* model using Huh7.5 cells.

**Figure 1:**
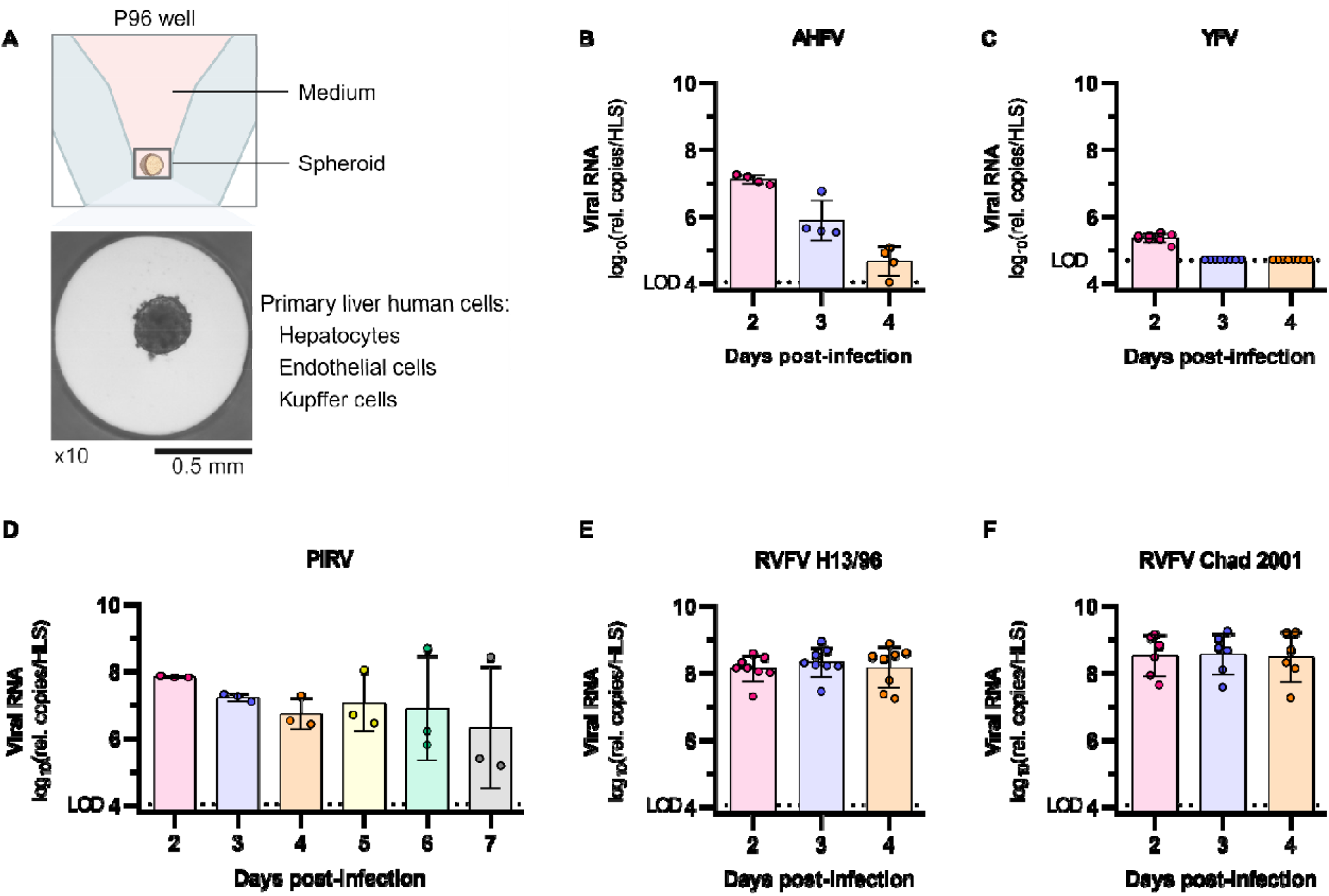
Infection of HLS with five BSL-3 HFVs. (A) Presentation of HLS model (created with BioRender.com). (B to F) Evaluation of the permissiveness of HLS to (B) AHFV, (C) YFV, (D) PIRV, (E) RVFV strain H13/96 and (F) RVFV strain Chad 2001, measuring viral RNA in cell supernatant with RT-qPCR on indicated dpi following an infection with a MOI of 10. Medium was completely renewed every day and measurement started on 2 dpi. Bars show mean ± standard deviation of value log_10_. Rel.: relative, LOD: limit of detection.

## 2. Materiel and methods

### 2.1. Experimental model

#### Cells and human liver spheroids (HLS)

African green monkey kidney cells (VeroE6, ATCC CRL-1586) were grown in minimal essential medium (MEM) supplemented with 7.5 % heat-inactivated foetal bovine serum (FBS), 1 % penicillin/streptomycin (P/S) and 1 % non-essential amino acids. Human hepatocellular carcinoma cells (Huh7.5, RRID CVCL_7927) were grown in Dulbecco’s Modified Eagle Medium (DMEM) high glucose and pyruvate supplemented with 10 % heat-inactivated FBS, 1 % P/S and 1 % non-essential amino acids (all medium and supplements are from ThermoFisher Scientific). Cells were maintained at 37□°C with 5 % CO_2_.

HLS (3D InSight™ Human Liver Microtissues, MT-02-302-04, InSphero) are composed of multi-donor primary human hepatocytes in co-culture with NPC, containing primary human Kupffer cells and primary liver endothelial cells (Fig. 1A). The NPC come from a unique donor and the hepatocytes from 10 different donors. They are cultivated and delivered in specific 96 well plates with approximately 1,000 cells per well. Once received, they were maintained at 37 °C with 5 % CO_2_ in 3D InSight™ human liver maintenance medium (TOX medium, CS-07-001-01, InSphero). Medium was changed every 2-3 days until the beginning of the experimentation.

#### Viruses

All experiments with infectious virus were conducted in a BSL-3 laboratory.

RVFV strain H13/96 (national collection of pathogenic viruses, NCPV 0310122v), RVFV strain Chad 2001 (French national reference center for Arboviruses, sequence currently being registered and available upon request), PIRV strain UVE/PIRV/UNK/VE/VAV488 (obtained through EVAglobal Evag Ref-SKU: 001V-02988, GenBank accession numbers: AF277659 et AY216505), AHFV strain 2010/EG/Shalatin2010-2_8S1 (obtained through EVAglobal, Evag Ref-SKU: 009V-03841, Genbank accession number: JX914664), YFV strain BOL 88/1999 (kindly provided by the National Centre of Tropical Diseases (CENETROP), Santa-Cruz, Bolivia, GenBank accession number: KF907504) were used in this study.

Virus stocks were prepared by inoculating a 25□cm^2^ culture flask of confluent VeroE6 or Huh7.5 (PIRV only) cells, respectively with MEM or DMEM medium supplemented with 2.5 % FBS and 1 % of P/S. The cell supernatant medium was replaced each 24□h and harvested at the peak of infection, supplemented with 25□mM HEPES (Thermofisher), aliquoted and stored at −80□°C.

### 2.2 Method details

#### Virus titration

To titrate viral productions and cell or spheroid supernatants, classical tissue culture infectious dose 50 % (TCID_50_) assays were performed on confluent VeroE6 cells in medium MEM + 2,5 % FBS + 1 % P/S. For that, 50 µL of ten-fold dilutions of cell or liver spheroid supernatants were used to inoculate cells in six replicates beginning respectively with 1/10 dilution for productions or 1/50 dilution for assays. After 7 days of incubation, the presence or absence of cytopathic effects was assessed microscopically for all viruses except PIRV. For this virus, a RT-qPCR was performed on the supernatant of each well as described below to determine if they showed viral replication. The infectious titers were estimated using the method described by Reed & Muench ^15^. The titer of PIRV production used for antiviral assays was determined using a focus forming assay performed on confluent VeroE6 cells in 96-well plate as follows: 100 µL of ten-fold dilutions of virus production in medium MEM + 2,5 % FBS + 1 % P/S were used to inoculate cells (six replicates per dilution). After an incubation at 37 °C with 5 % CO_2_ for 18 h, cells were fixed using 3,7 % paraformaldehyde (Sigma), then permeabilized and blocked with a solution of 0,1 % v/v Triton X-100 (Sigma) and 2 % w/v Bovine Albumin Serum (Sigma). To reveal infected cells, serum from immunized mice was used as primary antibody and a goat anti-mouse antibody (IgG, IgM (H+L) Secondary Antibody, Alexa Fluor™ 488, Thermofisher) as secondary antibody. Infected cells were counted through a fluorescent microscope (Invitrogen™ EVOS™ M5000 Imaging System). We used the replicates of the dilution showing 10 to 100 infected cells per well to calculate the viral concentration (focu-forming unit (ffu)/mL) as follows: mean number of infected cells * 10 * dilution factor.

#### Quantitative real-time RT-PCR (RT-qPCR) assays

To avoid contamination, all experiments were conducted in a molecular biology laboratory that is specifically designed for clinical diagnosis using molecular techniques, and which includes separate laboratories dedicated to perform each step of the procedure. Prior to PCR amplification, RNA extraction was performed using the QiaAmp 96 Virus, Qiacube HT kit and the Qiacube HT (both from Qiagen) following the manufacturer’s instructions. Shortly, 100 µL of supernatants or diluted supernatants were transferred into a S-block containing the recommended volumes of VXL, proteinase K and RNA carrier. RT-qPCR were performed with the GoTaq 1-step RT-qPCR kit (Promega) using 3.8 µL of extracted RNA and 6.2 µL of RT-qPCR mix that contains 250 nM of each primer and 75 nM of probe. Primers and probes sequences used are described in Table S1. Quantification was provided by four 2 log serial dilutions of an appropriate T7-generated synthetic RNA standard. Amplification was performed with the QuantStudio 12K Flex Real-Time PCR System (Applied Biosystems) using standard fast cycling parameters: 10 min at 50 °C, 2 min at 95 °C, and 40 amplification cycles (95 °C for 3 s followed by 30 s at 60 °C). Results were analysed using QuantStudio 12K Flex Applied Biosystems software v1.2.3.

#### Screening of virus in HLS

On day 0, after removing the medium from wells, liver spheroids were infected with a multiplicity of infection (MOI) of 10 using 70 µL of TOX medium. Each day, medium was renewed until day 4 post infection (dpi) or 7 dpi for PIRV. On the last day, 50 µL or 10 µL (YFV only) of spheroid supernatant were harvested and diluted respectively at 1/2 or 1/10 with Hanks’ balanced salt solution (HBSS, ThermoScientific) to perform RNA extraction followed by RT-qPCR as described above.

#### Characterization of viral replication in HLS

On day 0, after removing the medium from wells, HLS were infected with 70 µL of 3-fold serial virus dilutions starting from a MOI of 10 using TOX medium. Four wells were used as cell controls and were not infected. Each day, medium was renewed until 4 dpi (RVFV) or 7 dpi (PIRV). On each day from 2 dpi, 50 µL of spheroid supernatant were harvested and diluted at 1/2 with HBSS to perform RNA extraction followed by RT-qPCR as described above. For RVFV infection with strain Chad 2001, supernatants were used to perform a TCID_50_ as described above on 4 dpi.

On 4 dpi, cytopathic effects were measured using the CellTiter-Glo® 3D Cell Viability Assay (Promega) according to InSphero instructions. Briefly, reagent was prepared with 1:1 ratio with HBSS. The medium was removed from all wells and 40 µL of reagent were added per well. After mixing to lyse the spheroids, the content of each well was transferred to new plates to allow luminescence measurement. In empty wells, 40 µL of the same reagent were added to have well controls. Plates were incubated at room temperature on orbital shaking for 30 min. Luminescence (in counts per second (CPS)) was recorded for 1000 ms per well with a Tecan Infinite 200Pro machine (Tecan). The cell survival rates were calculated as follows: (CPS sample□−□CPS well control) / (CPS cell control□−□CPS well control).

#### Determination of RVFV infectivity in several models

In liver spheroids, virus infectivity was determined using values obtained from supernatant harvested on 4 dpi (MOI = 10; medium renewed every day as described above).

To determine the infectivity in Huh7.5 cell line, confluent cells in a 96 well culture plate were infected with a MOI of 1 in six wells and incubated for 2 days. After incubation, 100 µL of cell supernatant were used to do RNA extraction and RT-qPCR, and 5 µL were used to determine the infectious titer with TCID_50_ assay starting from 1/100 dilution, as described above.

To determine the infectivity *in vivo*, data from another study was used. *In vivo* experiments were approved by the local ethical committee (C2EA—14) and the French ‘Ministère de l’Enseignement Supérieur, de la Recherche et de l’Innovation’ (APAFIS#38977). Female C57BL6 mice (6 weeks) were provided by Charles River Laboratories. Animals were maintained in accordance with regulations. Isofluoran anaesthetized mice (Isoflurin®, Axience) were infected with 1,000 TCID_50_/mouse of RVFV (strain Chad 2001) in 100 µL of 0.9 % sodium chloride solution intraperitoneally. They were monitored daily to detect the appearance of any clinical signs of illness/suffering. Generally anaesthetized mice were euthanised by cervical dislocation on 1.5 dpi. A part of the liver median lobe was collected and weighted in a 2 mL tube containing 1 mL of 0.9 % sodium chloride solution and 3 mm glass beads. Samples were crushed and centrifuged prior to store supernatants at -80 °C. 100 µL of liver supernatant were used to do RNA extraction and RT-qPCR as mentioned before. 50 µL of cell supernatant were used to determine the infectious titers with a TCID_50_ assay as described above, starting from 1/10 dilution.

For all models, infectivity was calculated for each sample as follows: infectious titer (TCID_50_/mL of medium or g of liver) / viral load (relative copies/mL of medium or g of liver).

#### Half maximal effective concentration (EC_50_) and cytotoxic concentration (CC_50_) determination *in vitro*

One day prior to infection, 96 well culture plates (Greiner) were seeded with 3.6 x 10^4^ Huh7.5 cells in 100 µL assay medium per well (containing 2.5 % FBS and 1 % P/S). The next day, eight 2-fold serial dilutions of compounds at indicated concentrations (favipiravir (T-705) and nitazoxanide from BLDpharm; ribavirin and galidesivir (BCX4430) from MedChemExpress) in triplicate were added to the cells (25 µL/well, in assay medium). For the determination of the 50 % and 90 % effective concentrations (EC_50_, EC_90_; compound concentration required to inhibit by 50 % or 90 % viral RNA replication), eight “virus control” (VC) wells were supplemented with 25 µL of assay medium without any compounds. After 15□min, 25□µL of virus suspension, diluted in assay medium, was added to the wells with a MOI of 0.01 for RVFV and 1 for PIRV (except for cell controls). Eight “cell control” wells were supplemented with 50 µL of assay medium without any compounds or virus. Plates were incubated for 2 days at 37 °C prior to quantification of the viral genome by RT-qPCR as described above. For the determination of the 50 % cytotoxic concentrations (CC_50_; compound concentration required to reduce by 50 % cell viability), the same culture conditions were used, without addition of the virus, and cell viability was measured using CellTiter Blue® (Promega) following manufacturer’s instructions. Fluorescence (560/590□nm) was recorded with a Tecan Infinite 200Pro machine (Tecan).

Viral inhibition was calculated as follows: 100 * (quantity mean VC - sample quantity) / quantity mean VC. EC_50_, EC_90_ and CC_50_ were determined after performing a nonlinear regression (log(inhibitor) vs. response - Variable slope (four parameters)). All data obtained were analysed using GraphPad Prism 9 software (GraphPad software) ^16^. The selectivity index (SI) of the compounds was calculated as the ratio of the CC_50_ over the EC_50_.

#### Antiviral assay in HLS

On day 0, after removing the medium from wells, 20 µL of TOX medium were added to the wells. Then, HLS were treated with 25 µL of 3-fold serial dilutions of compounds in TOX medium in six replicates at indicated concentrations. As “virus control” (VC), wells were supplemented with 25 µL of TOX medium without compound. After 15□min, 25□µL of virus suspension diluted in TOX medium were added to the wells with a MOI of 10 (except for cell controls). “Cell control” wells were supplemented with 50 µL of TOX medium without any compounds or virus. Plates were incubated and medium was renewed, treating HLS as 0 dpi on each day. On 4 dpi, 10 µL of spheroid supernatant were harvested and diluted at 1/10 with HBSS to extract RNA and perform RT-qPCR. 10 µL of supernatants were also collected to do a TCID_50_ assay for RVFV. Cytopathic effects were measured using the CellTiter-Glo® 3D Cell Viability Assay. Viral inhibition was calculated as follows: 100 * (quantity geometric mean VC - sample quantity) / quantity geometric mean VC. Determination of % of cell survival rate were calculated as described above.

## 3. Results

### 3.1 RVFV and PIRV but not AHFV and YFV replicate in human liver spheroids (HLS)

We first evaluated the permissiveness of spheroid to several HFVs by infecting HLS with a high MOI (10). We assessed viral replication by measuring viral RNA by RT-qPCR in the culture medium, before fully renewing it daily (Fig. 1B to 1F). For AHFV, we observed an important decrease in the number of viral RNA copies from 2 to 4 dpi (Fig. 1B). This decrease over time appears to be linked to the gradual disappearance of viral RNA originating from the inoculum. In the case of YFV, only residual quantities of viral RNA were detected on 2 dpi (Fig. 1C). These results obtained with AHFV and YFV, and confirmed during a similar experiment (Table S2), indicate an inability to demonstrate viral replication for both viruses. Since PIRV replicates slowly in 2D cell models (data not shown), we decided to monitor the viral replication until 7 dpi (Fig. 1D). In two of the three wells infected with PIRV, we observed a gradual decline in viral RNA levels, suggesting a lack of viral replication. In contrast, viral replication was detected in one well, with viral RNA levels increasing from 7.3 log_10_ relative copies/HLS on 4 dpi to 8.7 log_10_ relative copies/HLS on 6 dpi. Regarding RVFV replication in HLS, for the laboratory strain H13/96 and clinical strain Chad 2001, significant viral RNA production was detected from 2 dpi, ranging from 7.3 to 9.1 log_10_ relative copies/HLS, and remained stable up to 4 dpi. This demonstrates that both RVFV strains were able to replicate in HLS.

### 3.2 Replication of PIRV and RVFV is dependent of MOI in HLS

We then decided to further explore the replication of PIRV and both strains of RVFV in this model. To determine if RVFV and PIRV replications are MOI dependent, we tested a range of six MOIs (Fig. 2). For that, HLS culturing media were analysed before renewing. We observed that PIRV replicated only at the highest MOI (3.33), for which the quantity of viral RNA peaked on 5 dpi, with an average of 7.5 log_10_ relative copies/HLS. However, as with a MOI of 10, significant variability was observed, with only two out of three wells showing viral replication (Fig. 2A). When infected with RVFV strain H13/96, viral replication was detected in all conditions: depending on the MOI, all or part of the wells infected on 4 dpi showed viral RNA production greater than 7 log_10_ relative copies/HLS (Fig. 2B). However, the less virus we inoculated, the more variability there was, with an increase in the number of uninfected HLS. Globally, the quantity of viral RNA increased from day to day in all conditions. For the clinical strain RVFV Chad 2001, the viral replication was also MOI-dependent. With the two highest MOI, 3.33 and 1.11, we measured respectively 8.2 and 7.9 log_10_ relative copies/HLS on 2 dpi and values remained globally constant on 3 and 4 dpi (Fig. 2C). With MOI of 0.04 and 0.01, the quantity of viral RNA increased from day to day with significant variability. For both strains of RVFV, results were confirmed by another experiments (Table S3).

**Figure 2:**
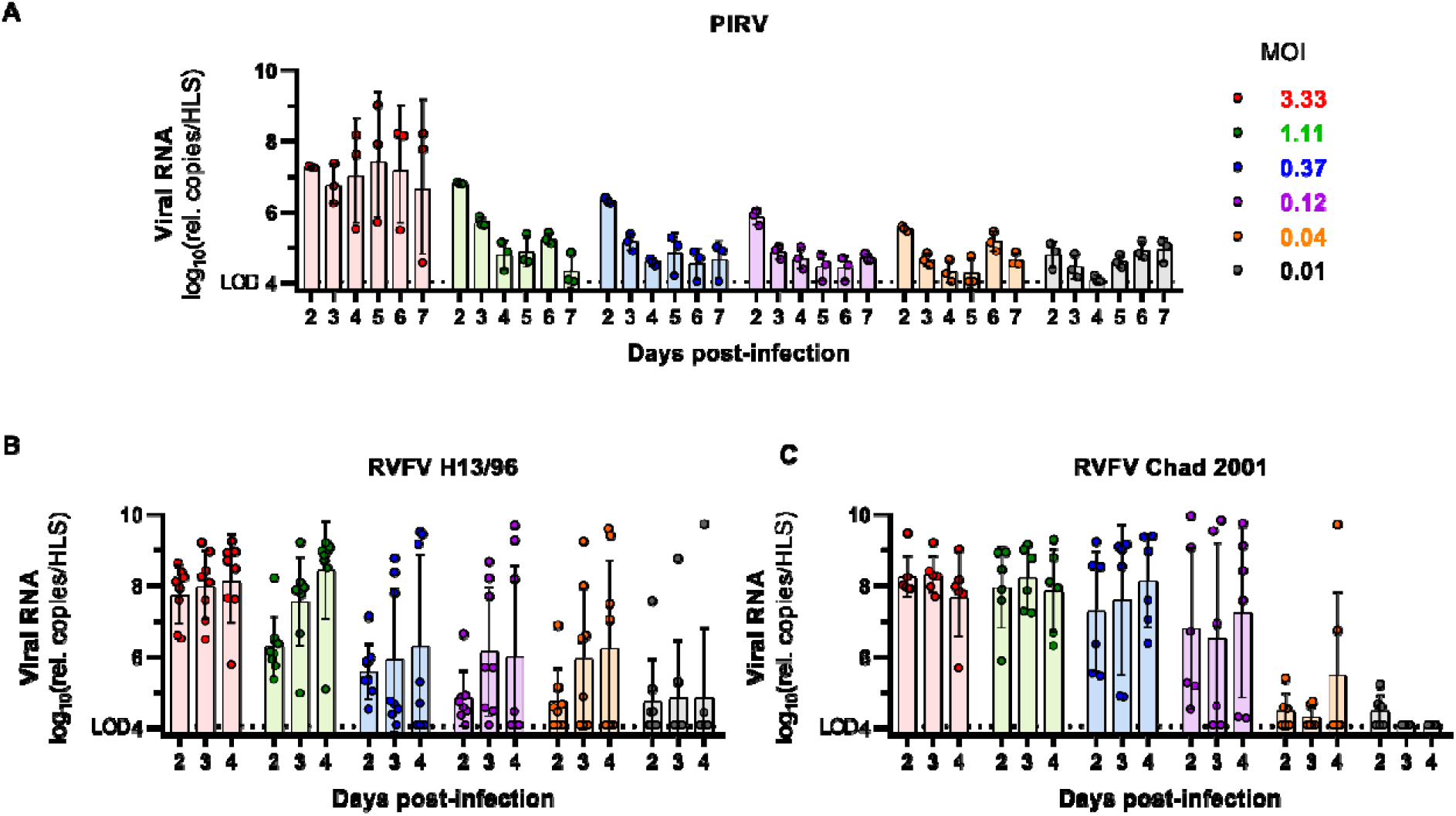
Replication kinetic of PIRV and RVFV with different MOI in HLS. Replication kinetic of (A) PIRV, (B) RVFV H13/96 and (C) RVFV Chad 2001 measuring viral RNA in cell supernatant with RT-qPCR on indicated days following an infection with indicated MOI. For all experiments, medium was renewed daily and measurement started on 2 dpi. Bars show mean ± standard deviation of value log_10_. Rel.: relative, LOD: limit of detection.

#### 3.3 RVFV Chad 2001 causes cytopathic effect in HLS and infectivity differs in comparison with other pre-clinical models

Consistently with growth characteristics in 2D cell culture, RVFV but not PIRV seemed to induce major cytopathic effect in HLS with a rupture of the spheroid integrity (Fig. 3A). To quantify this cytopathic effect, we performed a cell survival assay and observed a mean cell survival rate of 40 % on 4 dpi (Fig. 3B). To further characterise RVFV Chad 2001 infection in HLS, we measured the number of infectious particles and the quantity of viral RNA in the culture medium of HLS on 4 dpi (MOI of 10) (Fig. 3B). Results showed mean quantities of viral RNA of 9.7 log_10_ relative copies/HLS, which is similar with previous results (Fig. 1F), and mean infectious titers of 2.7 log_10_ TCID_50_/HLS. We then compared the relative infectivity (ie. infectious titer / relative viral RNA load) of the RVFV Chad 2001 in different preclinical models used for antiviral drug development: 2D cell culture of human hepatic cells (Huh7.5), a mouse model (liver samples) and HLS. HLS was the model with the lowest infectivity, showing values that were about 800-fold and 8-fold lower compared to Huh7.5 and mouse liver, respectively (Fig. 3C).

**Figure 3:**
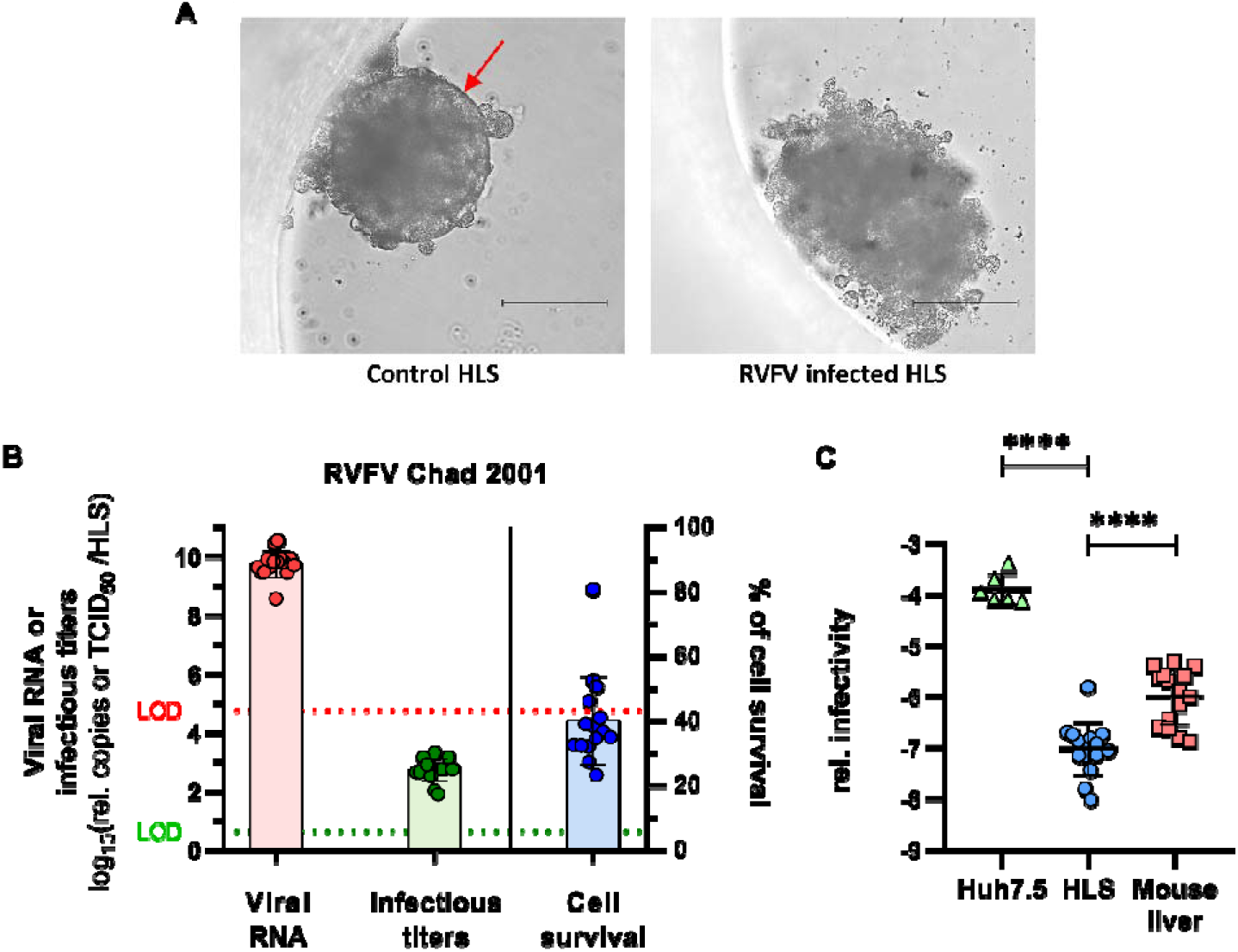
Characterisation of RVFV Chad 2001 infection in HLS. (A) Microscopic observation of HLS displaying control (left) or RVFV infected (right) with a MOI of 10 on 3 dpi (magnification x 20). The red arrow indicates the extracellular matrix. The scale bar indicates a length of 150 µm. (B) Characterisation of RVFV Chad 2001 infection in terms of released viral RNA copies (measured by RT-qPCR), infectious particles (TCID_50_ assay) and induced cytopathic effects on 4 dpi. (C) Infectivity of RVFV in several positive samples: Huh7.5 infected cells (n = 6), HLS (n = 15) or mouse liver (n = 15). Bars show mean ± standard deviation of value log_10_. Rel.: relative, LOD: limit of detection. ****: p-value < 0.0001 (Mann-Whitney test).

### 3.4 Inhibition of viral replication in HLS is more variable and requires higher compound dose in comparison with 2D cell line

To assess the suitability of HLS for antiviral assays, we evaluated the effect of broad-spectrum antivirals against PIRV and RVFV Chad 2001. For that, we selected favipiravir, galidesivir and ribavirin (only for RVFV), because of their known *in vitro* antiviral effect against these viruses or related ones ^17–19^. Nitazoxanide was added to this list for its *in vitro* broad-spectrum antiviral activity against various RNA viruses ^20–23^. We first confirmed the antiviral effect of these molecules and determined the effective dose of the compounds in Huh7.5 cells against both viruses (Table 1). Nitazoxanide and favipiravir had an EC_50_ around 0.5-1 µM against both viruses. Galidesivir showed EC_50_ of 33 µM and 18 µM against PIRV and RVFV respectively, and ribavirin has an EC_50_ of 26 µM against RVFV. All these results are coherent with previous studies ^17–19^.

**Table 1:** Activity of favipiravir, nitazoxanide, ribavirin and galidesivir against RVFV Chad 2001 and PIRV in Huh7.5 cells.

|  |  | EC<sub>50</sub> (µM) | EC<sub>90</sub> (µM) | CC<sub>50</sub> (µM) | Selectivity Index (SI) |
| --- | --- | --- | --- | --- | --- |
| PIRV | Nitazoxanide | 0.7 | 4.3 | 8.9 | 12.4 |
|  | Favipiravir | 1.1 | 1.5 | > 400 | > 367.0 |
|  | Galidesivir | 33.0 | 251.2 | > 100 | > 3.0 |
| RVFV Chad 2001 | Nitazoxanide | 0.6 | 1.3 | 8.9 | 14.6 |
|  | Favipiravir | 0.6 | 4.6 | > 400 | > 714.3 |
|  | Galidesivir | 17.7 | 37.6 | > 100 | > 5.6 |
|  | Ribavirin | 25.5 | 39.3 | 324.3 | 12.7 |

We then performed antiviral assays in HLS with the MOI that shown the highest replication associated with the lowest variability. Overall results showed dose-dependent effects of the molecules against both viruses (Fig. 4 and 5):

**Figure 4:**
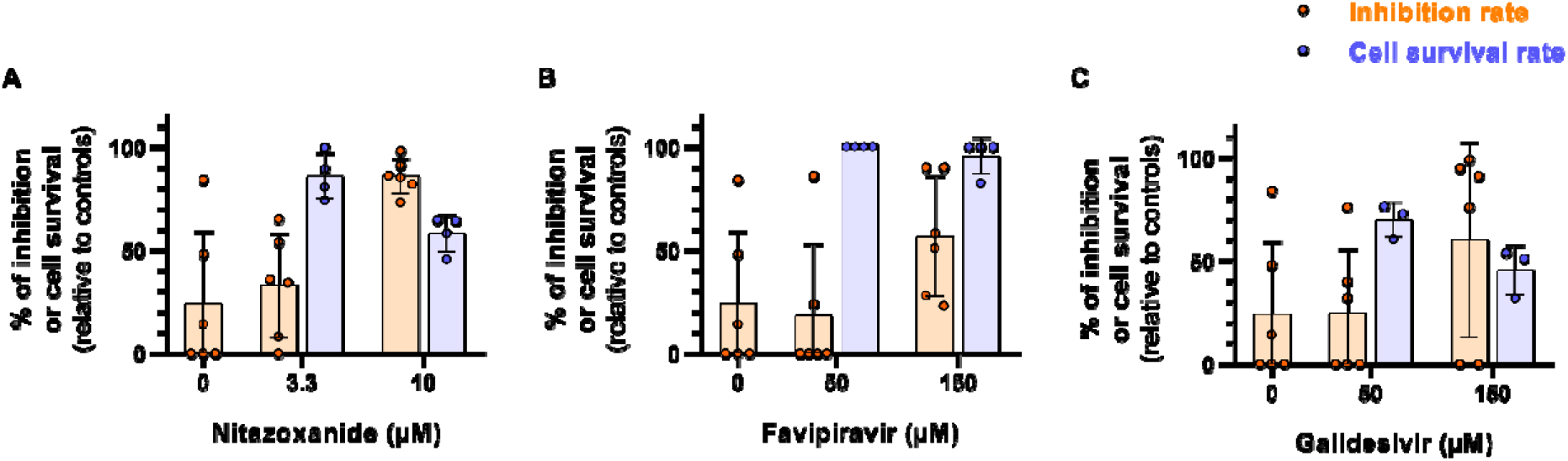
Activity of broad-spectrum antiviral molecules against PIRV in HLS. Antiviral activity of (A) nitazoxanide, (B) favipiravir and (C) galidesivir against PIRV in HLS. They were infected with a MOI of 10, medium and compounds were renewed every day, and measurements were made on 4 dpi. Percentages of inhibition and cell survival were obtained relative to control wells (untreated, called virus controls (VC) in methods section). Bars show mean ± standard deviation.

**Figure 5:**
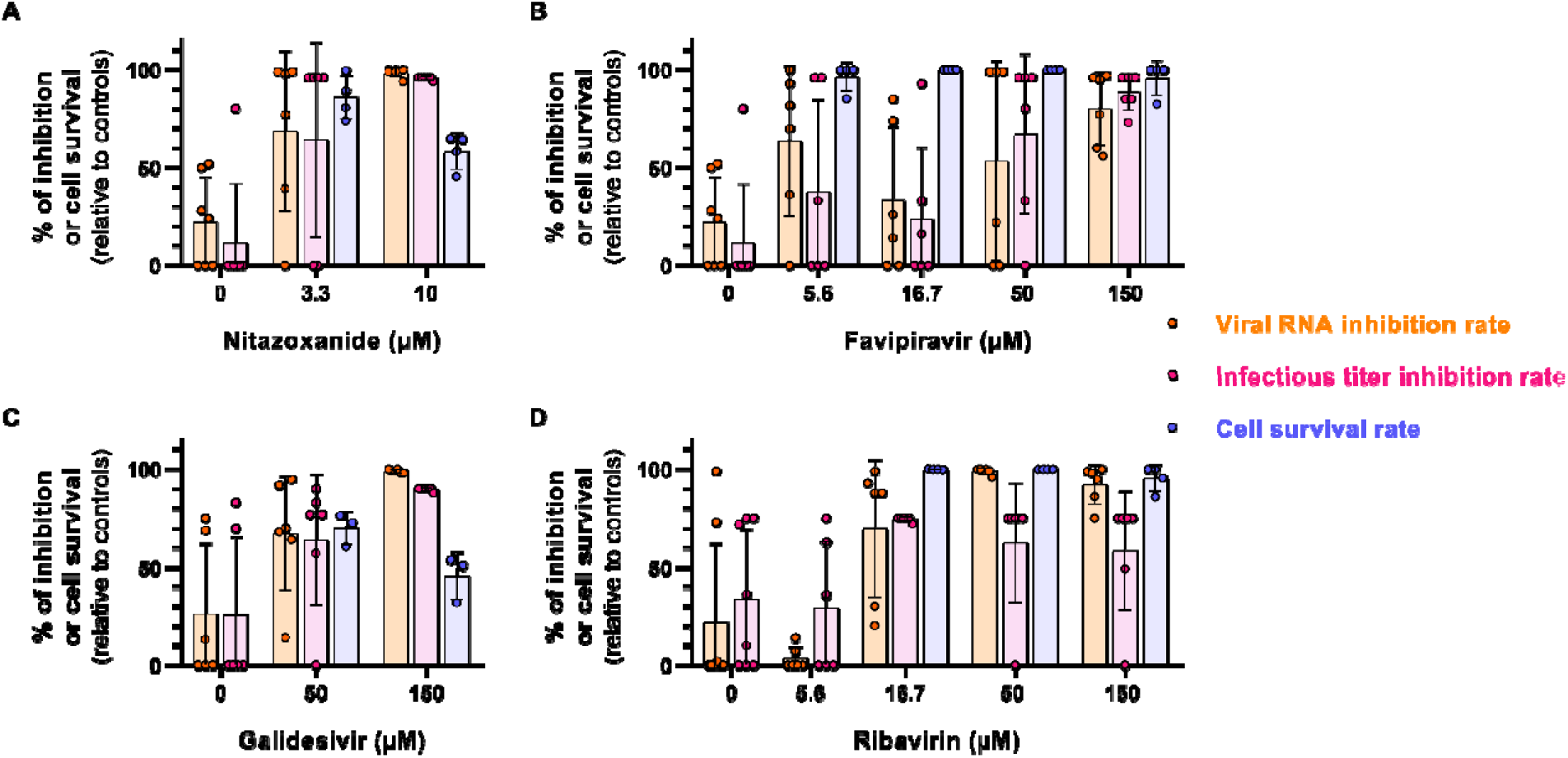
Activity of broad-spectrum antiviral molecules against RVFV Chad 2001 in HLS. Antiviral activity of (A) nitazoxanide, (B) favipiravir, (C) galidesivir and (D) ribavirin against RVFV Chad 2001 in HLS. They were infected with a MOI of 10, medium and compounds were renewed every day, and measurements were made on 4 dpi. Percentages of inhibition and cell survival were obtained relative to control wells (untreated, called virus controls (VC) in methods section). Bars show mean ± standard deviation.

#### Pirital virus

First, it is important to note that variability was observed in untreated wells (called virus virus control (VC) wells). At 3.3 µM, nitazoxanide inhibited viral RNA production by 33 % and the cell survival rate was 86 %. At 10 µM, cell survival rate dropped to 58 % and inhibition rate rose to 86 %, making it challenging to distinguish between toxicity and antiviral effect (Fig. 4A). Despite no cytotoxicity, favipiravir induced a limited antiviral effect with an RNA inhibition rate of 57 % at 150 µM, compared to the low EC_50_ and EC_90_ obtained in Huh7.5 cells (Table 1 and Fig. 4B). Such as nitazoxanide, galidesivir induced cytotoxic effects, with cell survival rates of 70 % and 46 % at 50 and 150 µM respectively. The corresponding inhibition rates of viral replication were highly variable (25 % and 60 %, respectively) (Fig. 4C). All the values used to calculate the percentages of inhibition are available in Table S4.

#### Rift Valley fever virus

Given that RVFV is cytopathic in 2D cell lines, we also performed a TCID_50_ assay on culture media to estimate the inhibition rate of infectious titer, complementing the analysis based on viral RNA production (Fig. 5). As with PIRV, untreated wells exhibited important variability. Despite this, antiviral assays against RVFV in HLS showed a more efficient inhibition of viral replication than for PIRV, with inhibition rates of viral RNA yield above 80 % for all compounds at the highest concentrations tested. HLS treated with 3.3 µM of nitazoxanide, a concentration associated with high cell survival rate (86 %), showed an inhibition of viral replication rate of around 70 % (Fig. 5A). As with PIRV, the inhibition rates were very high (> 95 %) at 10 µM of nitazoxanide but significant toxicity was also observed at this concentration. With favipiravir, we observed a dose-dependent effect on viral replication, but the compound concentration required to inhibit 50 % of viral replication was more than 50 times higher than the EC_50_ measured in Huh7.5 cells (Fig. 5B, Table 1). Results showed an inhibition of viral replication rate of around 30 % at 16.7 µM and above 80 % at 150 µM. The results for nitazoxanide and favipiravir were confirmed with one supplemental experiment (Fig. S1 and Table S5). Treating HLS with 150 µM of galidesivir induced inhibition rates above 90 % (Fig. 5C). However, as described for PIRV, galidesivir is cytotoxic in HLS with 45 % of cell survival at 150 µM making difficult to demonstrate an antiviral effect. Finally, ribavirin which is not cytotoxic in HLS at all tested concentrations, showed a consistent dose-dependent antiviral effect from 16.7 µM with inhibition rates around 70 % at this concentration. All the values used to calculate the percentages of inhibition are available in Table S6.

## 4. Discussion

The handling of haemorrhagic fever viruses (HFVs) is strictly regulated, potentially limiting the speed of development of specific research models. *In vitro* 2D cell models are the easiest and most cost-effective models to implement in high biosecurity level laboratories. However, they lack the functions and interactions necessary to faithfully reflect the complexity of the human body. *In vivo* models are obviously the most accurate way of mimicking human pathophysiology, but they are time-consuming to implement and costly. Organoids have been increasingly developed recently, and represent promising tools for studying a wide range of functionalities, as well as being complementary to 2D cell models, in particular to reduce the need for *in viv*o models in accordance with the 3R rule ^12^. Therefore, it is important to carefully evaluate some of them, particularly for the purposes of HFVs drug development. In this study, we evaluated a ready-to-use HLS model, as HFVs typically exhibit hepatic tropism. Each HLS is composed of around 1,000 primary human cells, including multi-donor hepatocytes and single donor of NPC.

In the first part of this study, we assessed the permissiveness of HLS to a panel of BSL-3 HFVs. To this end, we successfully established a procedure for the simultaneous quantification of viral RNA production (RT-qPCR), infectious titers (TCID_50_ assay, for RVFV only) in the supernatant of HLS, all in a 96-well format. To complete the assessment of viral replication, we also performed a cytotoxicity assay adapted for 3D cell cultures, to evaluate viral cytopathicity and compound cytotoxicity. Previous studies have demonstrated that these models can also be utilized for cytokine and fluorescence assays ^24,25^. Thus, using such a relevant model could enhance our capacity to better understand the pathophysiology of viral infections and enable us to study antiviral compound mechanisms of action or attenuated strains.

We first attempted to infect HLS with two orthoflaviviruses, AHFV and YFV. We found that both viruses did not replicate in this model. In mice, AHFV replicates only in Kupffer cells and not in hepatocytes ^26,27^. Immunocompromised mice exhibit a high level of AHFV replication in the liver and symptoms correlated with the most severe human cases ^27^. However, liver involvement in immunocompetent mice or pigtailed macaques is less significant reflecting results obtained in HLS ^26,28^. In contrast to AHFV, YFV is well known to infect hepatocytes and Kupffer cells, inducing necrosis of hepatocytes preferentially in the midzonal part of the liver ^29–32^. The first explanation could be that YFV did not replicate in HLS because the cell receptors involved in YFV infection are absent from the HLS surface. Another possible explanation could be a donor-dependent protection against YFV, as only a small proportion of humans infected with YFV develop hepatic involvement. Since NPC come from a unique donor and hepatocytes from 10 different donors, the number of susceptible cells could be insufficient to allow robust viral infection and replication. This donor-dependent protection might be natural or acquired through prior vaccination against YFV. Another study showed that the inter-human variability of Kupffer cells within a similar organoid model had an impact on toxicological responses after exposure to nanomaterials, with differences observed in cytokine release ^25^. Moreover, it has been demonstrated that monocytes and macrophages, such as Kupffer cells, can be “trained” from previous infections, leading to a more efficient response to new infections through epigenetic and transcriptomic modifications ^33–36^.

We then tested two viruses that belong to the *Hareavirales* order, PIRV, used as a surrogate for new-world BSL-4 mammarenaviruses and the phlebovirus RVFV. Despite some variability, PIRV and RVFV appeared to replicate in HLS and these replications were dependent of MOI. These findings are consistent with *in vivo* observations, where both viruses are known to infect hepatocytes and to a lesser extent, Kupffer cells (endothelial cells are infected only by PIRV) ^37–42^.

With a lower MOI, we were able to detect replication only for RVFV, and not in all the HLS tested. This notable feature is conserved *in vivo*, with the need for higher doses of PIRV to infect adult laboratory hamsters (10^4^ pfu/hamster) compared to RVFV (10 pfu/hamster) ^38,39,43^. With both viruses, the high MOI-dependent infection and variability may be linked to the fact that the primary cells may have retained their ability to express antiviral genes. Indeed, it has been shown that cells were able to activate this antiviral response more effectively with a low MOI than a high MOI ^44^.

We compared the infectivity of the clinical strain of RVFV in different models used for drug development: 2D cell culture (Huh7.5 cells), HLS and a mouse model. Our results revealed that the virus replicated in the hepatic 2D cell line with higher efficiency resulting in virions with higher infectivity compared to *in vivo*, in the liver of mice and in HLS. These results highlighted that replication and viral production are cell-dependent and not very efficient in HLS. However, the fact that infectivity in HLS is more similar to that in mouse liver suggests that this model more closely reflects *in vivo* conditions.

We then assessed in HLS the antiviral activity of some already well-described antiviral compounds against PIRV and RVFV, to evaluate whether this model could bridge the gap between 2D cell lines and *in vivo* studies for antiviral development. Once again, we were confronted with great variability in viral replication, possibly due to the difficulties of having strong, and robust replication in HLS. Despite this variability, the analysis of antiviral activity for both viruses indicated that higher concentrations of favipiravir, galidesivir, and nitazoxanide were required to inhibit viral replication, compared to the results observed in Huh7.5 cells. The use of a high MOI to infect HLS represents a first hypothesis to explain this difference. The most significant difference in the inhibition rate of viral RNA production between 2D cell lines and HLS was observed for favipiravir. This compound demonstrated an antiviral effect by protecting hamsters and rats from succumbing to RVFV infection at relatively low doses ^45,46^. Similar results were obtained against Lassa and Pichinde viruses, two mammarenaviruses showing similar EC_50_ than PIRV in 2D cell culture models ^19,47,48^. This discrepancy between the activity of favipiravir in HLS and other models could be explained by the fact that favipiravir is a prodrug that becomes active after intracellular phosphoribosylation, which converts it into its active form, favipiravir-RTP, which inhibits the RNA-dependent RNA polymerase of RNA viruses ^49^. This phosphoribosylation has been documented to be cell-dependent ^50^ which could explain the difference observed between antiviral activity in 2D cell lines and that in HLS. Our results also revealed a cytotoxic effect of nitazoxanide and galidesivir at tested concentrations. Regarding the nitazoxanide, its toxicity was similar to that observed in Huh7.5 cells. Galidesivir, on the other hand, appears to be more toxic in HLS than in Huh7.5 cells. Its toxicity was also observed in hamsters, despite the absence of serious adverse effects and the overall safety of the compound when administered to healthy adult subjects ^17,51^. In the hamster model, galidesivir demonstrated antiviral efficacy against RVFV, although this efficacy was limited by its toxicity, similar to that observed in HLS ^17^.

In conclusion, although a high MOI was required to infect HLS and some variability in viral replication was observed, our results suggest that HLS are susceptible to infection by viruses belonging to the order *Hareavirales*, unlike orthoflaviviruses. Moreover, we were able to evaluate antiviral activity and cytotoxicity effect of known antivirals against PIRV and RVFV. However, results obtained for antiviral assays with favipiravir do not reflect 2D cell line and *in vivo* results. Altogether, our results suggests that HLS could be used to obtain relevant information that complements data from 2D cell culture models but not as a systematic selection step during pre-clinical evaluation of antiviral molecules. Furthermore, such model could be useful to study the barrier of resistance and the mechanism of action of antiviral compounds, as well as their safety, since liver is a key organ for drug metabolism ^52^. In a more general way, HLS could be considered to study mechanisms of replication, underlying factors of viral pathogenicity or host-pathogen interactions.

## Acknowledgements

We thank Camille Placidi and Géraldine Piorkowski (UVE; Marseille) for their valuable technical contribution. A part of the work was done on the Aix Marseille University antivirals platform “Plateforme Criblage Viral Marseille-Timone (PCVMT)” belonging to the Marseille Screening Center “MaSC”. We also thank the National Center of Reference for Arboviruses for kindly providing the clinical RVFV strain.

S.C. is the recipient of a thesis scholarship funded by the French Ministry of Defence - Defence Innovation Agency and Aix-Marseille University. F.T. is supported by the IRD Chair “Antiviral strategy for emergence in the South”, in partnership with Aix-Marseille University, Inserm and ANRS MIE. This project is funded by the French Agence Innovation Defense (FHeTALS project (ANR-20-ASTR-0009)). The activity of the UVE is supported by its supervising institutional bodies (Aix-Marseille Université, Università di Corsica, Institut national de la santé et de la recherche médicale, Institut de recherche pour le développement, Institut de Recherche Biomédicale des Armées). This project was supported by the ANRS-MIE (PRI projects of the EMERGEN research program).

## Author contributions

Conceptualization, S.C., J.-S.D., A.N. and F.T.; Methodology, S.C., J.-S.D., A.N. and F.T.; Validation, S.C., J.-S.D., A.N. and F.T.; Formal Analysis, S.C. and F.T.; Investigation, S.C. and F.T.; Resources, X.d.L, A.N. and F.T.; Writing-Original Draft, S.C., J.-S.D., A.N. and F.T.; Writing – Review & Editing, S.C., J.-S.D., A.N. and F.T.; Visualization, S.C., J.-S.D., X.d.L., A.N. and F.T.; Supervision, X.d.L., A.N. and F.T.; Funding acquisition, A.N.

## Supplemental Data

**Table S1**: RT-qPCR systems. Related to Methods.

**Table S2**: Confirmation of the lack of permissiveness of human liver spheroids (HLS) to Alkhurma haemorrhagic fever virus (AHFV) and yellow fever virus (YFV).

**Table S3**: Confirmation of the replication kinetic of Rift Valley fever virus (RVFV) strains H13/96 and Chad 2001 with different MOI in HLS.

**Table S4**: Values used to calculate the percentages of inhibition of viral RNA or infectious particles production of Pirital virus (PIRV).

**Figure S1**: Confirmation of the activity of nitazoxanide (A) and favipiravir (B) against RVFV Chad 2001 in HLS.

**Table S5**: Values used to calculate the percentages of inhibition of viral RNA or infectious particles production of RVFV Chad 2001.

**Table S6**: Values used to calculate the percentages of inhibition of viral RNA or infectious particles production of RVFV Chad 2001.

